# Transfer Learning for Survival-based Clustering of Predictors with an Application to *TP53* Mutation Annotation

**DOI:** 10.1101/2025.10.06.680732

**Authors:** Xiaoqian Liu, Hao Yan, Haoming Shi, Emilie Montellier, Eric C. Chi, Pierre Hainaut, Wenyi Wang

## Abstract

*TP53* is the most frequently mutated gene in human cancers, and germline mutations in *TP53* cause Li-Fraumeni syndrome (LFS), a hereditary predisposition to diverse cancers. Accurate annotation of *TP53* mutations based on their survival effects is critical for informed LFS patient management. Motivated by this need, we develop a new approach for Survival-based Clustering of Predictors (SCP) by identifying homogeneous coefficients in Cox regression. We formulate this task as a fusion-penalized Cox regression problem and provide an efficient computational algorithm. A nonconvex distance-to-set penalty is adopted to facilitate parameter tuning and improve estimation accuracy. To overcome data limitations, we further develop TLSCP, a transfer learning extension that borrows coefficient ranking information from a source dataset under the assumption of similar ranking patterns between source and target. TL-SCP integrates ranking information through weighted rank averaging, allowing flexibility in accommodating cohort heterogeneity while maintaining model simplicity. Simulation studies demonstrate TL-SCP’s superior performance over SCP in clustering recovery and coefficient estimation. In the application of *TP53* mutation annotation where we utilize non-LFS germline *TP53* mutation carriers as a source cohort for the target LFS cohort, TL-SCP identifies biologically meaningful *TP53* mutation clusters and offers improved clinical interpretability compared to experiment-based annotations.

## 1 Introduction

The practical problem that motivates this work is the annotation of *TP53* mutations. Known as the “guardian of the genome”, the *TP53* gene plays a critical role in cell signaling, apoptosis, metabolism, DNA repair and transcription (Levine, 2021). At the same time, it is also the most frequently mutated gene in human cancers (Vogelstein et al., 2000), underscoring the high clinical relevance of *TP53* mutations. In particular, germline *TP53* mutations (inherited and present in all cells) are the primary cause of Li-Fraumeni syndrome (LFS), a rare hereditary cancer predisposition that features a high lifetime risk (up to 90%) for a broad spectrum of cancers including early-onset and even childhood cancers (Li and Fraumeni Jr, 1969). Moreover, affected individuals face an elevated risk of subsequent primary cancers as well as treatment-related secondary cancers. It is widely acknowledged that the wide functional gradient of different germiline *TP53* mutations contributes to LFS heterogeneity (Olivier et al., 2009; Rocca et al., 2022). Nevertheless, it remains poorly understood how different *TP53* mutations relate to various phenotypic consequences and survival outcomes in LFS, which is clinically crucial for personalized treatment and genetic counseling.

The *TP53* community has devoted tremendous efforts in understanding such relationships and annotating the *TP53* mutational landscape. Most studies in the literature have focused on individual mutations, particularly hotspot mutations that are frequently observed in LFS patients, such as R175H (Chiang et al., 2021) and R337H (Galante et al., 2025). One pioneering initiative is the ClinGene *TP53* Variant Curation Expert Panel (VCEP, Fortuno et al. (2021)) which brings together experts from various fields to group clinically relevant *TP53* variants associated with LFS and establish standardized guidelines for clinical variant interpretation. More recently, research has shifted toward data-driven approaches for systematic annotation of *TP53* mutations. For instance, some studies (Montellier et al., 2024; Fischer et al., 2025) classify *TP53* variants based on their biological functions, leveraging experimental measures such as transcriptional activity scores (Kato et al., 2003) or cellular functional screens (Giacomelli et al., 2018). Phenotypic differences in LFS are then investigated across the resulting functional variant clusters.

While experiment-based annotation of *TP53* mutations may serve as reference for clinical interpretation, a substantial gap remains between experimentally characterized functions of *TP53* and observed LFS phenotypes. We hypothesize that directly leveraging patient outcomes for mutation annotation offers a more clinically grounded perspective with greater translational relevance for advancing patient management. To the best of our knowledge, no prior studies have investigated *TP53* mutation annotation through the lens of patient survival data. Here, we introduce the first statistical framework to cluster *TP53* mutations based on time-to-event clinical outcomes, while simultaneously addressing the fundamental challenge posed by data limitations in LFS studies.

Specifically, to link various *TP53* mutations with LFS patient survival outcomes, we treat mutations as survival predictors in the Cox proportional hazards model. Our goal is to annotate *TP53* mutations based on their survival effects, quantified by the Cox regression coefficients. By assuming that mutations share common or similar effects in their unknown clusters, we recast the mutation annotation problem as identifying homogeneity among the Cox regression coefficients. To this end, we build on the homogeneity pursuit strategy (Ke et al., 2015) and develop Survival-based Clustering of Predictors (SCP), a fusion-penalized Cox regression approach that clusters predictors into groups with homogeneous effects. We present a general computational framework for implementing SCP that flexibly accommodates different penalty choices. In this work, we particularly employ a nonconvex distance-to-set penalty (Chi et al., 2014; Xu et al., 2017), which enables direct specification of the number of clusters while improving estimation accuracy.

A major obstacle for survival analysis of LFS and *TP53* is the limited availability of data. Germline pathogenic *TP53* mutations are rare, with an estimated prevalence of approximately 1 in 3, 000 to 1 in 10, 000 individuals in the general population (de Andrade et al., 2024). The prevalence of LFS is even lower, resulting in a limited number of well-documented cases. Moreover, *TP53* mutations predominantly occur at some hotspots, with ten variants accounting for roughly 30% of cases, while the majority of individual mutations remain poorly characterized with scarce carriers (Stiewe and Haran, 2018). Collectively, these limitations pose significant challenges for the reliable application of statistical methods, including the proposed SCP, in *TP53* mutation annotation. To mitigate this, we leverage transfer learning, an emerging technique for overcoming data limitations.

The core idea of transfer learning is to improve learning performance on a target cohort with limited sample size by borrowing information from one or more larger, related source cohorts. Stemming from computer science, transfer learning has found widespread applications in customer review classification (Pan and Yang, 2009), natural language processing (Alyafeai et al., 2020), and genetic risk prediction (Lu et al., 2024), to name a few. Recently, transfer learning has gained growing attentions from statistics and been investigated in various statistical scenarios, including high-dimensional linear regression (Li et al., 2022), generalized linear models (Tian and Feng, 2023), Gaussian graphical models (Li et al., 2023a), and large-scale quantile regression (Jin et al., 2024). Some recent studies (Li et al., 2023b; Xie et al., 2024) have further extended transfer learning to Cox regression for time-to-event outcomes, operating under the assumption that the source and target tasks share similarity in the magnitude of their regression coefficients.

Inspired by these developments, we adapt transfer learning to our proposed SCP method but in a distinct manner. Specifically, we transfer coefficient ranking information from the source to the target, under the assumption that ranking patterns remain similar across the two tasks. In our motivating application, it is biologically plausible to assume different germline *TP53* mutations share a similar ranking of survival significance between the non-LFS *TP53* mutation carriers and the LFS patients, where the former cohort finds a larger sample size in the National Cancer Institute (NCI) *TP53* Database (de Andrade et al., 2022). In the proposed SCP framework, a preliminary ranking of the Cox regression coefficients is required, enabling the transfer of informative ranking patterns to improve clustering accuracy. We refer to the transfer learning-enhanced SCP method as TL-SCP, in which ranking transfer is implemented via weighted rank averaging. TL-SCP is built upon the same methodological backbone as SCP, yet offers practical advantages in data-limited settings. Of note, although motivated by the problem of *TP53* mutation annotation, the proposed SCP and TL-SCP are broadly applicable to clustering survival predictors in general settings, with TL-SCP additionally requiring a source dataset.

In summary, this work makes the following contributions:

1. We propose an SCP approach to cluster survival predictors by searching for homogeneous coefficients in Cox regression using fusion penalization. We provide an efficient computational framework for SCP with various options of penalties, among which the distance-to-set penalty is advocated for its practical merits.
2. To resolve the data limitation issue, we further develop the TL-SCP method, which innovatively transfers the coefficient ranking information from the source data to improve the model performance on the target data. This TL-SCP method enjoys high flexibility in accommodating cohort heterogeneity while maintaining model simplicity for practical consideration.
3. We are the first to annotate *TP53* mutations by their survival effects on LFS patient outcomes. Our study using the NCI *TP53* Database suggests improved clustering performance using TL-SCP. Our clustering result displays clearer survival differences across mutation groups and offers better clinical variant interpretation, compared to existing experiment-based *TP53* annotations.

The rest of the paper is organized as follows. Section 2 first introduces the basic SCP method with its computational algorithm and then develops the TL-SCP extension with details of the involved transfer learning. Section 3 evaluates both SCP and TLSCP through comprehensive simulation studies. Section 4 presents the analysis of *TP53* mutation annotation and its validation. Section 5 concludes with a discussion.

## 2 Method

### 2.1 The SCP method

Consider the survival data given in the form of 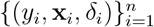, where the observed time *y*_*i*_ is a time of failure if *δ*_*i*_ = 1 or right-censoring if the indicator *δ*_*i*_ = 0, and 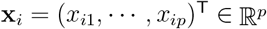 is the vector of *p* predictors (or covariate variables). Let *t*_1_ *< t*_2_ *< · · · < t*_*m*_ be the increasing sequence of unique failure times, and *j*(*i*) denotes the index of the observation failing at time *t*_*i*_. The Cox proportional hazards model (Cox, 1972) assumes the following semi-parametric hazard function

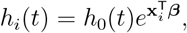

where *h*_*i*_(*t*) is the hazard for patient *i* at time *t, h*_0_(*t*) is a shared baseline hazard, and ***β*** *∈* ℝ^*p*^ is the vector of coefficients quantifying the survival effects of the *p* predictors. Then the partial likelihood can be written as

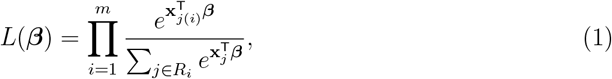

where *R*_*i*_ = *{j*: *y*_*j*_ *≥ t*_*i*_*}* is the set of indices of patients at risk at time *t*_*i*_. Note that the partial likelihood (1) assumes unique *y*_*i*_’s, but it can be suitably modified for the case of ties as discussed in Simon et al. (2011). When ***β*** possesses a certain structure, for example ***β*** is sparse, estimation of ***β*** is typically achieved by minimizing the negative log-partial likelihood augmented with a penalty function Ψ(***β***) as follows:

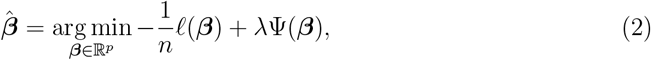

where the log-partial likelihood *l*(***β***) is expressed as

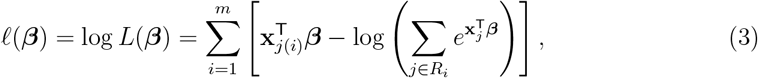

*λ >* 0 is the regularization parameter, and the penalty function Ψ(***β***) enforces the desired structure in ***β***. For instance, the Lasso penalty Ψ(***β***) = *∥****β****∥*_1_ imposes sparsity in ***β*** (Tibshirani, 1996).

For homogeneous survival effects, *β*_*i*_’s share common values in their unknown clusters. Then homogeneity pursuit represents simultaneous coefficient estimation and cluster recovery. Motivated by the homogeneity pursuit strategy (Ke et al., 2015) for linear regression, we extend it to Cox regression and summarize the procedure as follows:

1. Obtain a preliminary estimator 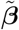 from the classical Cox model.
2. Construct the rank mapping *{τ* (*j*): 1 *≤ j ≤ p}* such that 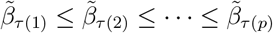.
3. Compute the estimate by solving a fusion penalized Cox regression problem (2) with

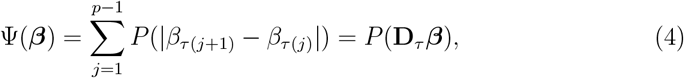

where *P* is a sparsity-inducing penalty that shrinks some pairwise coefficient differences towards zero, thereby identifying clusters of predictors with shared coefficients. **D**_*τ*_ *∈* ℝ ^(*p−*1)*×p*^ is a fusion matrix constructed according to the rank mapping *τ*, and its *i*-th row is defined as **e**_*τ*(*i*)_ *−* **e**_*τ*(*i*+1)_ where **e**_*j*_ *∈*ℝ^*p*^ denotes the standard basis vector with the *j*-th element to be 1.

We refer to the aforementioned framework as Survival-based Clustering of Predictors (SCP). Alternative choices for the penalty function Ψ(***β***) include the total variation penalty (Harchaoui and Lévy-Leduc, 2010) and a hybrid pairwise penalty proposed in Ke et al. (2015), both of which can be expressed as a fusion penalty of the form Ψ(***β***) = *P* (**D*β***). The primary difference among these different penalties lies in the fusion matrix **D**, which determines the specific coefficient pairs subjected to the penalty *P* on their differences. For SCP, we opt for the simplest fusion penalty (4) that only consider adjacent pairs (after ranking) for two key reasons: 1) it is computationally efficient and 2) it establishes a direct relationship between the sparsity level (i.e., the number of nonzero elements in **D**_*τ*_ ***β***) and the number of clusters in ***β***, which we will discuss shortly. Indeed, the success of SCP relies on the consistency between *τ* and the ranking of the true coefficients ***β***\*. With this in mind, we use transfer learning to enhance preliminary rankings (Section 2.3).

The way of constructing the fusion matrix **D**_*τ*_ in the SCP framework implies that the number of clusters in ***β*** equals the number of nonzeros in **D**_*τ*_ ***β*** plus one. In practice, users often prefer tuning the number of clusters directly. Traditional sparsity-inducing penalties, such as Lasso (Tibshirani, 1996), SCAD (Fan and Li, 2001), and MCP (Zhang, 2010), are not satisfying in this regard. This motivates us to introduce the distance-to-set penalty to serve as the sparsity-inducing penalty *P* to make SCP more user-friendly.

To estimate a parameter vector ***θ*** subject to a set constraint ***θ*** *∈ C*, the distance-to-set penalty (Chi et al., 2014; Xu et al., 2017) is defined as

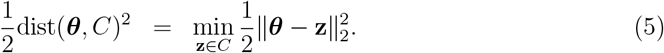

To achieve sparsity, one can define *C* = *{***z**: *∥***z***∥*_0_ *≤ k}* where *k* is the desired number of nonzero elements in **z**. A family of proximal distance algorithms has been developed for solving the distance-penalized problem min_***θ***_ 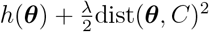 for different types of loss functions *h*(***θ***) (Keys et al., 2019; Landeros et al., 2022a). In practice, *λ* is iteratively sent to *∞*, so the effective tuning parameter is *k*, allowing direct control over sparsity. Distance penalization has been successfully applied in various statistical learning settings (Landeros et al., 2022b; Liu et al., 2023; Landeros et al., 2025), demonstrating superior estimation accuracy and support recovery while facilitating parameter tuning.

By employing the distance-to-set penalty in (4), SCP solves the following distance fusion-penalized Cox regression problem

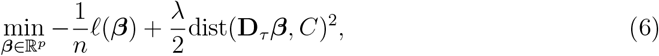

where the sparsity level *k* corresponds to the number of nonzeros in **D**_*τ*_ ***β***. Given the explicit relation between *k* and the number of clusters in ***β***, users are able to directly tune the number of clusters. Solving problem (6) can be handled at ease by the computational framework we proposed for SCP, as described in the next section.

### 2.2 Algorithm for SCP

To compute the SCP estimate, the key is to solve the penalized Cox regression problem (2) with Ψ(***β***) = *P* (**D**_*τ*_ ***β***). Following Simon et al. (2011), we approximate the log-partial likelihood (3) via a quadratic expansion, hence transforming the problem into iteratively solving a sequence of fusion-penalized weighted least squares problems (7). The complete computational framework for SCP is summarized in Algorithm 1. For detailed technical derivations, we refer readers to Simon et al. (2011).

A key advantage of Algorithm 1 is its ‘plug-and-play’ nature: any fusion-penalized least squares solvers can be incorporated to implement SCP with the corresponding fusion penalty. This flexibility allows us to readily solve the distance fusion-penalized Cox model (6) using existing proximal distance algorithms (Landeros et al., 2022a). In our SCP implementation, options for the sparsity-inducing penalty *P* include the Lasso, SCAD, MCP, and the distance-to-set penalty.

### 2.3 The TL-SCP method

Our motivation for incorporating transfer learning into the SCP framework comes from both theoretical and practical considerations. In theory, SCP requires the preliminary estimate 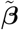 to have a ranking consistent with the true ***β***\*, which in practice demands a sufficiently large sample size to obtain a reliable preliminary estimate. On the contrary, clinical data is often limited, as exemplified in our *TP53* application. While adopting more complicated penalties, such as the total variation penalty, could theoretically relax the ranking consistency requirement, in practice they incur high computational cost and often suffer from model selection issues, especially with small sample sizes. Therefore, we retain the simple fusion penalty (4) for SCP to leverage its practical advantages (Section 2.1) and turn to transfer learning to address data limitations.

#### Algorithm 1

**Computational framework for SCP**

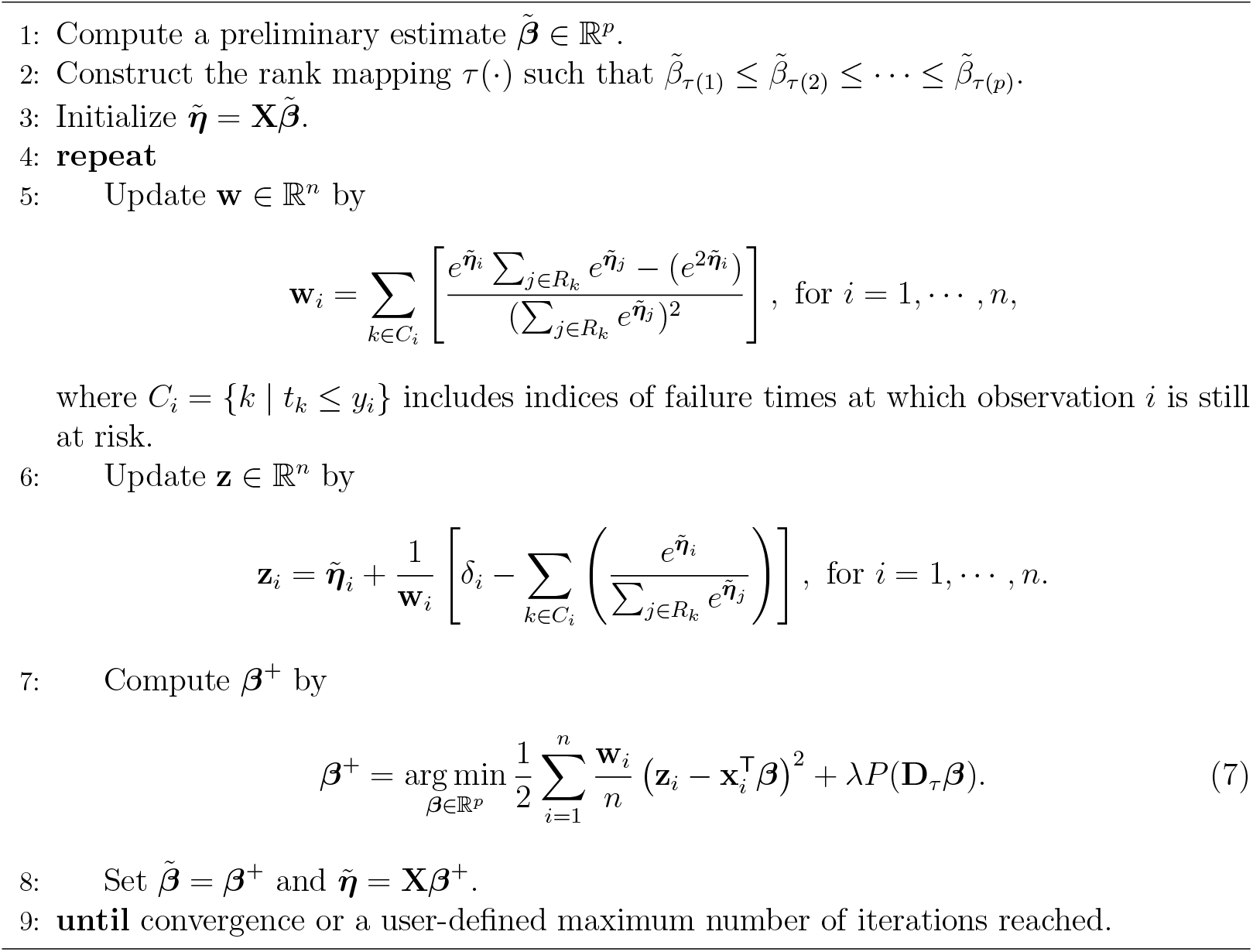

Consider a target cohort 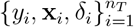 and a source cohort 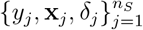with *n*_*T*_ and *n*_*S*_ independent observations, respectively. Here by default, both cohorts share the same set of predictors **x** = (*x*_1_. *· · ·*, *x*_*p*_)^T^. In general, we assume *n*_*S*_ is much larger than both and *p*. Given cohort *c ∈ {T, S}*, the hazard function reads

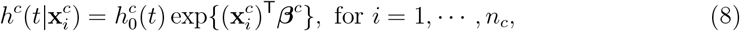

where 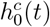 is the cohort-specific baseline hazard function, and ***β***^*c*^ is the cohort-specific coefficient vector. When relying solely on the target cohort, the estimation of ***β***^*T*^ is hindered by the limited sample size *n*_*T*_. Transfer learning improves estimation by leveraging source information, typically under the assumption that ***β***^*T*^ and ***β***^*S*^ are similar in magnitude, enforced via sparsity on their difference ***β***^*T*^ *−* ***β***^*S*^ (Li et al., 2023b; Xie et al., 2024).

#### Algorithm 2

**Computational framework for TL-SCP**

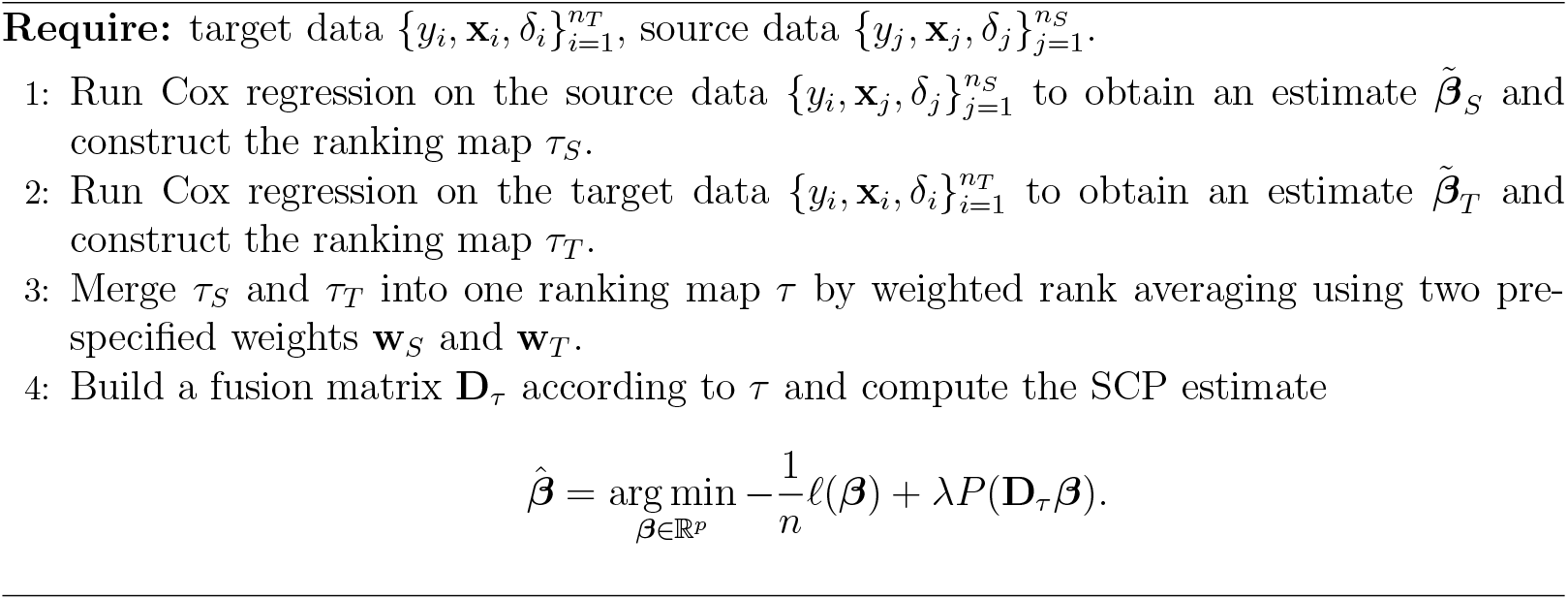

This work adopts transfer learning for Cox regression in a novel manner. In settings with a limited target cohort, the bottleneck of SCP is obtaining a reliable preliminary ranking of the regression coefficients ***β***^*T*^. To address this, we assume that ***β***^*T*^ and ***β***^*S*^ share a similar ranking structure and develop the transfer learning SCP (TL-SCP) method to effectively borrow ranking information from the source cohort.

TL-SCP operates as follows. First, we fit the classical Cox regression models separately on the source and target cohorts and obtain the respective ranking maps *τ*_*S*_ and *τ*_*T*_ from the estimated coefficients. Next, we construct a unified ranking map *τ* by weighted rank averaging with pre-specified weights **w**_*S*_ and **w**_*T*_. For example, we may set 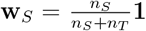 and 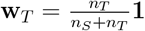, where **1** denotes a vector of ones. Finally, we use the weighted ranking *τ* to build a fusion matrix **D**_*τ*_ for SCP to cluster predictors. The complete procedure for TL-SCP is presented in Algorithm 2. Of note, when data-sharing is restricted, TL-SCP can simply ask for the estimated coefficients 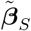 or the ranking map *τ*_*S*_ from the source, avoiding the need for individual-level data.

We conclude this section by emphasizing the flexibility and generalizability of TL-SCP. First, the assumed ranking similarity is less restrictive than the conventional magnitude similarity typically imposed in transfer learning frameworks, and is therefore more likely to hold in practice. Second, potential differences in predictor distributions between cohorts, known as covariate shift in transfer learning, are accommodated by fitting Cox models separately for source and target cohorts rather than directly pooling the data. Third, the weighted rank averaging step integrates ranking information from both cohorts and flexibly controls their contributions via tunable weights, thus adjusting for potential ranking discrepancies. Forth, the TL-SCP framework can be naturally extended to multiple source scenarios. It can also be generalized to cases where the target predictor set is a subset of the source predictor set, provided that the ranking similarity assumption holds for overlapping predictors. Last but not the least, TL-SCP brings little burden in the computational cost compared to running SCP solely on the target cohort, as fitting the source Cox model and performing weighted rank averaging are both computationally cheap.

## 3 Numerical Studies

We conducted extensive simulation studies to demonstrate the efficacy of our proposed SCP method and the improved performance of TL-SCP in identifying clusters of predictors with homogeneous survival effects. With a source cohort available, we compared three methods: 1) SCP using the target cohort only, 2) TL-SCP using both the source and target cohorts, and 3) a naive baseline method by running the standard Cox regression on the target and then using k-means to cluster the obtained coefficients, denoted as ‘Cox-kmeans’. Although ‘Cox-kmeans’ is not able to produce homogeneous coefficient estimates, it can serve as a naive approach for clustering predictors based on their survival effects, thereby included in our simulation studies as a baseline for clustering performance comparison. For both SCP and TL-SCP, we employed the distance-to-set penalty as illustrated in (6). We additionally conducted a series of simulation examples to demonstrate the outperformance of SCP with the distance-to-set penalty in comparison to SCP with commonly used penalties including Lasso, SCAD and MCP. For space consideration, we relegated these simulations on penalty comparisons under various scenarios to the supplementary material.

### 3.1 Simulation settings

#### Data generation

We fixed the target cohort and varied the data generation of the source cohort to mimic different real-world scenarios. For the target cohort, the sample size *n*_*T*_ was set to 200. We generated the *i*-th observed vector of predictors **x**_*i*_ *∈* ℝ^*p*^ with *p* = 90 by sampling *x*_*ij*_ i.i.d. from a uniform distribution on [*−*0.5, 0.5]. To generate homogeneous coefficients, we first set the true coefficient vector 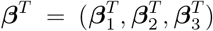, where each 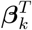 was a constant vector of length *p/*3, with entries taking common values 2, 0.5, and *−*1, respectively. We then randomly permuted the elements of ***β***^*T*^ to hide the cluster structure in ***β***^*T*^. The survival time *y*_*i*_ of the *i*-th individual was then generated from the exponential distribution with the rate parameter 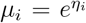, where 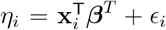 and *ϵ*_*i*_ *∼ N* (0, 0.5^2^) is the Gaussian random noise. For the censoring indicator vector 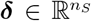, we specified the censoring rate *r* to be 0.3 and then randomly selected *rn*_*S*_ elements from *{*1, *· · ·, n*_*S*_*}* to be the censoring indices, i.e., the corresponding *δ*_*i*_ = 0. For the source cohort, its data generation varied according to different source scenarios and will be described specifically under each scenario in Section 3.2.

### Tuning parameter selection

For both SCP and TL-SCP methods, the tuning parameter is the integer *k* associated with the distance-to-set penalty. We employed the Bayesian Information Criterion (BIC) to select *k* from a candidate set *{*1, 2, 3, 4, 5*}*. For TL-SCP, we set the weight parameters in the weighted rank average step to be the corresponding sample size ratios. For the baseline ‘Cox-kmeans’ method, we used the average silhouette width approach to determine the number of clusters in k-means.

### Performance evaluation

We evaluated the compared methods on their performance in both coefficient estimation and cluster recovery. Specifically, we used the relative squared error, defined as 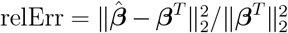, to quantify the estimation accuracy. We used the normalized mutual information (NMI, Ana and Jain (2003)) to measure the similarity between the recovered clustering result and the ground truth. Let *𝒞* = *{C*_1_, *C*_2_, *· · ·}* and *𝒟* = *{D*_1_, *D*_2_, *· · ·}* be two sets of disjoint clusters of *{*1, *· · ·, p}*, the NMI is defined as

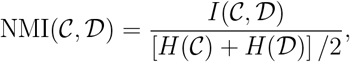

where *I*(*𝒞, 𝒟*) =ℑ *i*,*j* (|*𝒞*_*i*_ *∩𝒟* _*j*_|*/p*) log(*p*|*C*_*i*_ *∩𝒟* _*j*_|*/*|*𝒞*_*i*_||*𝒟* _*j*_|) denotes the mutual information between *𝒞* and *𝒟*, and *H*(*𝒞*) = *−* ℑ_*i*_(*𝒞*_*i*_*/p*) log(|*C*_*i*_|*/p*) denotes the entropy of *𝒞*. The NMI takes values on [0, 1], and a larger NMI value implies a higher similarity between two clustering results.

To reflect complex real-world scenarios, we conducted various simulation experiments under different source scenarios regarding the source sample size, the covariate shift, and the ranking inconsistency between the source and target coefficients. For each simulation experiment, we ran 50 replicates for each method and used boxplots to report the median as well as the 25th and 75th quantiles of the two performance metrics.

### 3.2 Simulations under different source scenarios

#### Consistent ranking without covariate shift

We started with the simplest scenario where covariate variables (predictors) in both source and target cohorts shared the same distribution, namely, there was no covariate shift. In addition, we assumed the ranking of coefficients was consistent between the two cohorts. Specifically, for the source cohort with a given sample size *n*_*S*_, its *i*-th observed vector of predictors were generated from a uniform distribution on [*−*0.5, 0.5] as in the target cohort. To ensure consistency in the coefficient ranking between two cohorts, we set ***β***^*S*^ to be the log transformation of the coefficient ranking of the target by setting ***β***^*S*^ = log(rank(***β***^*T*^, ties.method = “first”)) in R. The generation of **y** and ***δ*** for the source cohort and the associated parameters remained the same as in the target cohort. We varied the sample size *n*_*S*_ of the source cohort and summarized the simulation results of the compared methods in Figure 1.

**Figure 1.**
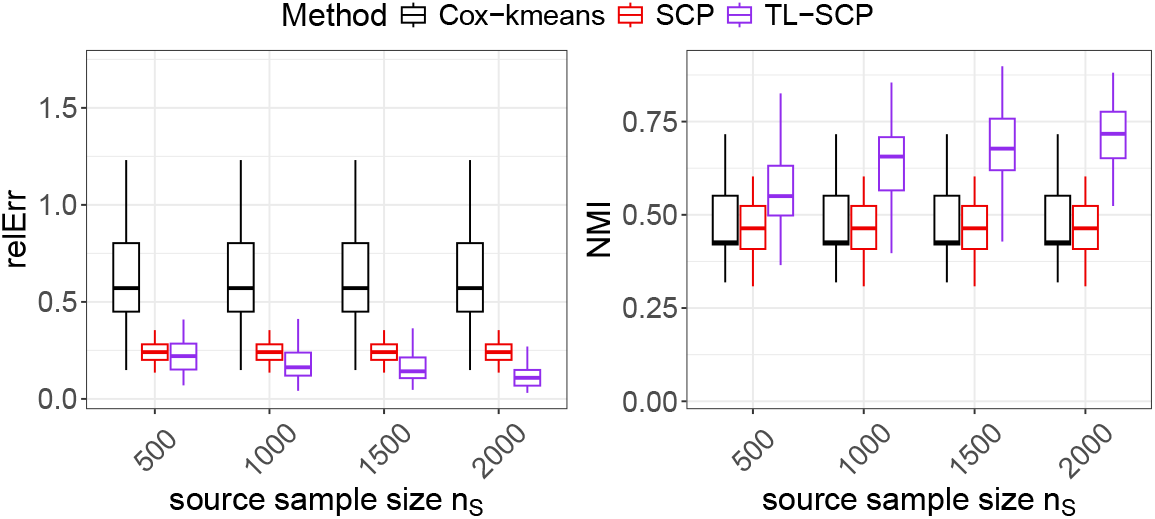
Performance comparison of TL-SCP to SCP and Cox-kmenas under the scenario of ‘consistent ranking without covariate shift’.

As shown in Figure 1, when using the target cohort only, SCP achieved notable lower estimation errors and higher NMI values than the naive Cox-kmeans approach, suggesting its superior performance in both coefficient estimation and cluster recovery. However, the median NMI values of both SCP and Cox-kmeans were below 0.5, highlighting the inherent challenge of achieving accurate clustering using only the target dataset. With the availability of a source cohort, TL-SCP demonstrated improved performance over SCP in both coefficient estimation and cluster recovery. In particular, when the source sample size exceeded 1000, TL-SCP achieved median NMI values above 0.7, indicating strong cluster recovery performance.

#### Consistent ranking with covariate shift

To examine the effect of covariate shift on the performance of TL-SCP, we modified the generation of predictor vectors in the source cohort. Given a sample size *n*_*S*_ = 1000 and number of predictors *p* = 90, we generated the source predictor vector **x**_*i*_ by setting *x*_*ij*_ = *z*_*ij*_ + *ζ*_*ij*_ where 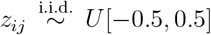 and 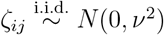 for *i* = 1, *· · ·, n*_*S*_ and *j* = 1, *· · ·, p*. That is, we added a covariate shift by adding a normal variable *ζ*_*ij*_ and used its standard deviation *ν* as a representative of the shift level. The generation of the source coefficient ***β***^*S*^ remained the same as in the first experiment, so that ranking consistency was guaranteed. We applied the same procedure to generate **y** and ***δ*** for the source cohort as in the target cohort. We varied the shift level *ν* from 0.05 to 0.5. The simulation results under different values of *ν* were shown in Figure 2.

**Figure 2.**
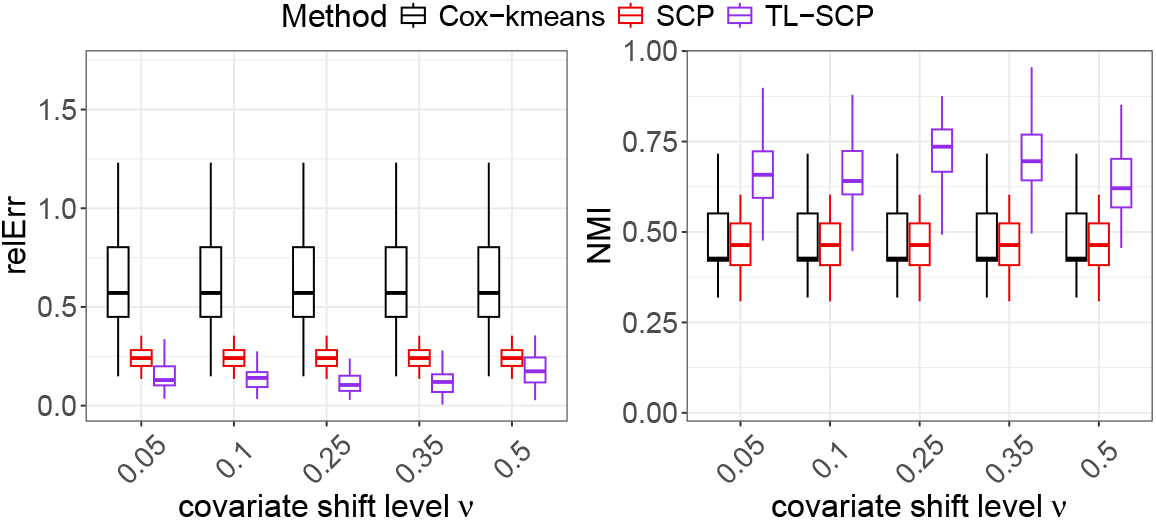
Performance comparison of TL-SCP to SCP and Cox-kmenas under the scenario of ‘consistent ranking with covariate shift’.

As Figure 2 displays, TL-SCP consistently outperformed SCP across varying levels of covariate shift, exhibiting lower estimation errors and higher NMI scores. Under a substantial shift *ν* = 0.5 where the covariate shift nearly overrides the baseline, TL-SCP maintained clearly superior clustering results while achieving estimation accuracy comparable with SCP. In addition, TL-SCP’s performance remained stable as *ν* increased, with no clear signs of degradation. These results highlight the robustness of TL-SCP to the covariate shift between the source and target cohorts.

### Inconsistent ranking with covariate shift

A major practical concern of using TL-SCP would be the potential inconsistency of coefficient ranking between the source and target cohorts. To address this concern, we further test the ability of TL-SCP in accommodating inconsistent coefficient rankings between the two cohorts. For the source data generation, we fixed the sample size *n*_*S*_ = 2000 and the number of predictors *p* = 90 and simulated the source predictors following the procedure in the second experiment with a fixed shift level *ν* = 0.25. Regarding simulating the source coefficients, we introduced a parameter *s* to quantify the ranking inconsistency level between the source and target coefficients. Specifically, after generating the target coefficients ***β***^*T*^ as described in Section 3.1, we permuted the order of the first *s* elements of ***β***^*T*^ and then used the permuted ranking to generate the source coefficients ***β***^*S*^ by the log transformation as described in the first experiment. We varied the ranking inconsistency level by setting *s* = *{*5, 10, 15, 20*}*.

Figure 3 displayed the produced estimation errors and NMI scores of TL-SCP at different values of *s*. As anticipated, as the ranking inconsistency level *s* grew, the performance of TL-SCP degraded in both estimation accuracy and cluster recovery. Under the settings where more than 10 out of the 90 predictors had different coefficient rankings across the two cohorts, the median estimation errors of TL-SCP can be larger than those of SCP, but its median NMI scores remained higher. We reasoned that these high NMI scores were attributed to the weighted rank averaging step in TL-SCP, which utilized the majority of coefficients with consistent ranking to contribute to the improved clustering results.

**Figure 3.**
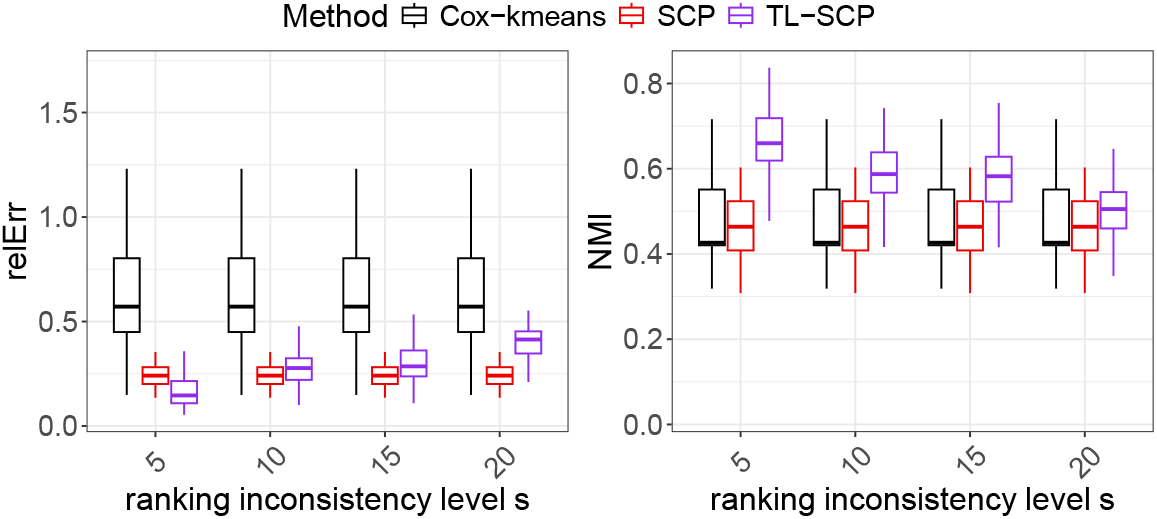
Performance comparison of TL-SCP to SCP and Cox-kmenas under the scenario of ‘inconsistent ranking with covariate shift’.

Together, the above simulation studies demonstrated that by effectively leveraging ranking information from the source cohort, TL-SCP substantially improved model performance on the target cohort in terms of both coefficient estimation and clustering recovery. Notably, these gains in clustering recovery were evident not only in the overall NMI metric but also in the detailed clustering assignments. To illustrate this, the supplement presents contingency tables for a representative example across different methods. In brief, Cox-kmeans produced only two clusters, as the cox model itself does not yield homogeneous estimates and kmeans is unable to differentiate them. SCP produced five clusters, with three major clusters aligning with the true clusters. TL-SCP identified three clusters, with minor discrepancies relative to the true labels. Further details of this example were provided in the supplement.

## 4 *TP53* mutation annotation

We now apply the proposed SCP and TL-SCP methods to our motivating problem of *TP53* mutation annotation. The objective of our study was to annotate germline *TP53* mutations by their survival effects on LFS patient survival outcomes to provide enhanced clinical interpretability. In particular, the survival outcome of interest was the age at first cancer diagnosis, which has direct implications on choosing appropriate cancer screening regimes for individuals with germline *TP53* mutations (Kratz et al., 2017).

Our analysis was performed using the *TP53* Database, the largest publicly available *TP53* data resource hosted by the National Cancer Institute (NCI) of the United States (de Andrade et al., 2022). The dataset of germline *TP53* variants contains information on individuals that are carriers of a germline *TP53* mutation and families in which at least one family member has been identified as a germline *TP53* mutation carrier. Patients were annotated as LFS (Li-Fraumeni syndrome), LFL (Li-Fraumeni like), FH (family history of cancer), and no-FH (no family history of cancer). In general, LFS annotations reflect clinician-evaluated cancer patients, whereas LFL, FH, and no-FH primarily represent population-based collections of germline *TP53* mutation carriers. For our analysis, we therefore separated the dataset into LFS versus non-LFS groups, corresponding to clinically ascertained versus population-based cohorts. After removing individuals with missing mutation or age information, we obtained a cohort of 273 LFS patients with 46 confirmed recurring germline *TP53* mutations. Among these 46 mutations, 24 of them were associated with only two or three LFS patients, posing great challenges in accurate survival analysis. Since only a small proportion of germline *TP53* mutation carriers develops LFS, this motivated us to transfer the survival information from the larger population of general germline *TP53* mutation carriers (non-LFS patients) to assist the study of LFS. Through the same data pre-processing, we collected a cohort of 1484 germline *TP53* mutation carriers with 303 recurring *TP53* mutations, 39 of which also appeared in the LFS patient cohort. Therefore, we focused on the 39 overlapped mutations shared by the two cohorts, and the associated numbers of individuals were 255 LFS patients and 689 germline *TP53* mutation carriers, respectively. We denoted the 255 LFS patients as the target cohort and the 689 general germline *TP53* mutation carriers as the source cohort.

We applied TL-SCP to cluster the considered 39 germline *TP53* mutations by utilizing both the source and target cohorts and compared its performance with SCP applied solely to the target cohort. We treated mutations as survival predictors and included gender as a covariate. We employed the distance-to-set penalty for both TL-SCP and SCP given its superior performance observed in our simulation studies. For TL-SCP, we set the two weights for rank averaging as 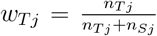 and 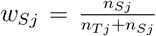, where *nT j* and *n*_*Sj*_ are the numbers of the *j*-th *TP53* mutation carriers in the corresponding cohorts. We again used the BIC for selecting the number of clusters from a candidate set *{*1, 2, 3, 4, 5*}* for each method. Both SCP and TL-SCP identified three mutation clusters, but their clustering structures differed. For ease of exposition, we labeled the three mutation groups with decreasing estimated coefficients as ‘early-onset’, ‘mid-onset’, and ‘late-onset’, reflecting their respective effects on age at first cancer diagnosis. SCP produced group sizes of 24, 14, and 1, with the singleton ‘late-onset’ cluster raising concerns about the reliability of the clustering. In contrast, TL-SCP produced more balanced group sizes of 23, 12, and 4.

We used an alluvial plot in Figure 4 to illustrate annotation discrepancies between SCP and TL-SCP, with individual mutations listed for each TL-SCP group and mutations with discrepant annotations highlighted in color. A closer inspection of individual mutations further supported the validity of the TL-SCP clustering. For example, common hotspot mutations such as R175H, R248W, R273H, and R282W were clustered in the ‘early-onset’ group, consistent with findings from relevant studies (Klemke et al., 2021; Sun et al., 2020; Zhang et al., 2016). Additionally, R337C, strongly associated with LFS (Davison et al., 1998) and classified as pathogenic by the VCEP, was labeled ‘early-onset’ by TL-SCP, whereas SCP annotated it as ‘mid-onset’. The splice variant T125T was classified into the TL-SCP ‘mid-onset’ group but was labeled as ‘early-onset’ by SCP. This discrepancy was driven by the two LFS patients in the target cohort with early-onset cancers (ages 2 and 27). In contrast, the source cohort included an additional 30 T125T mutation carriers exhibiting a broad range of ages at cancer onset. By leveraging this information, TLSCP corrected the annotation to ‘mid-onset’, yielding a classification more consistent with clinical observations (Pinto et al., 2022). TL-SCP annotated R213Q and R267W as ‘lateonset’, which aligned with their ability in retaining transactivation activities (Petitjean et al., 2007; Pan and Haines, 2000), whereas SCP labeled them as ‘mid-onset’. A complete comparison of individual mutation annotations from SCP and TL-SCP was summarized in Table B.1 and allocated to the Supplement.

**Figure 4.**
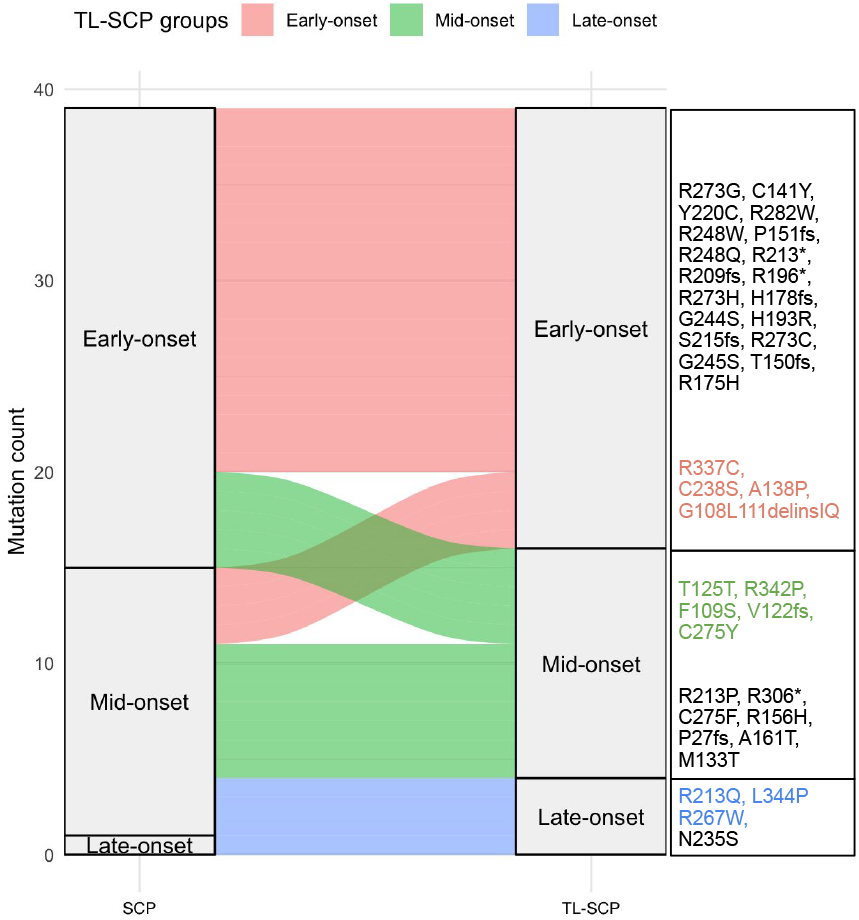
Alluvial plot of *TP53* mutation annotations (SCP vs. TL-SCP). The right-hand text boxes list individual mutations in each TL-SCP group. Mutations highlighted in color indicate those with discrepant annotations between SCP and TL-SCP.

To further evaluate the biological utility of the clustering result from TL-SCP, we compared a *TP53* mutation grouping system using the mutations’ functional activities, which were quantified in yeast-based transactivation assays (YTA) (Montellier et al., 2024). These assay-based measures remain popular to serve as differential scores for missense *TP53* mutations, even though researchers are constantly looking for new approaches to improve beyond this system (Hoyos et al., 2022; Montellier et al., 2025). Using the hierarchical Ward’s clustering method on eight YTA scores, Montellier et al. (2024) obtained four functional missense mutation classes, named by ‘A’, ‘B’, ‘C’ and ‘D’, and further included a class of ‘0’ to denote nonsense and frame-shift mutations. These five mutation classes formed three mutation groups exhibiting distinct survival patterns in age at first cancer diagnosis, as demonstrated in their analysis relating YTA classes to LFS clinical phenotypes. Specifically, classes ‘0’ and ‘A’ showed similar, most severe cancer-onset profiles, so we combined them into an ‘early-onset’ group. YTA class ‘B’ displayed an intermediate onset pattern, hence we call it the ‘mid-onset’ group. YTA classes ‘C’ and ‘D’ were associated with attenuated onset profiles, hence we name it the ‘late-onset’ group. Among the 39 *TP53* mutations in our study, 33 were labeled as ‘early-onset’ by the YTA method, 3 as ‘mid-onset’, and 2 as ‘late-onset’, resulting in a highly unbalanced clustering structure. In addition, the splice variant T125T was not examined in Montellier et al. (2024) and therefore lacked a YTA annotation. An alluvial plot in Figure 5 compared annotation results between YTA and TL-SCP, and a detailed comparison of individual mutation annotations was provided in Table B.1 in the Supplement. We observed that all 23 TL-SCP ‘early-onset’ mutations were also classified as ‘early-onset’ by YTA, demonstrating strong concordance between the two approaches, while TL-SCP further consolidated the remaining mutations into ‘mid-onset’ and ‘late-onset’ groups, yielding a more balanced clustering structure. Furthermore, LFS patients stratified by the three TL-SCP mutation groups exhibited distinct and well-separated survival patterns in age at first cancer diagnosis (right panel of Figure 6), providing additional validation of the clustering results. In contrast, survival patterns for the YTA ‘mid-onset’ and ‘late-onset’ groups showed considerable overlap, particularly for patients with age below 30 (left panel of Figure 6). This comparison demonstrated that the more balanced clustering from TL-SCP is more informative for annotating *TP53* mutations according to their effects on age at first cancer diagnosis, which could offer valuable clinical guidance for LFS patient management.

**Figure 5.**
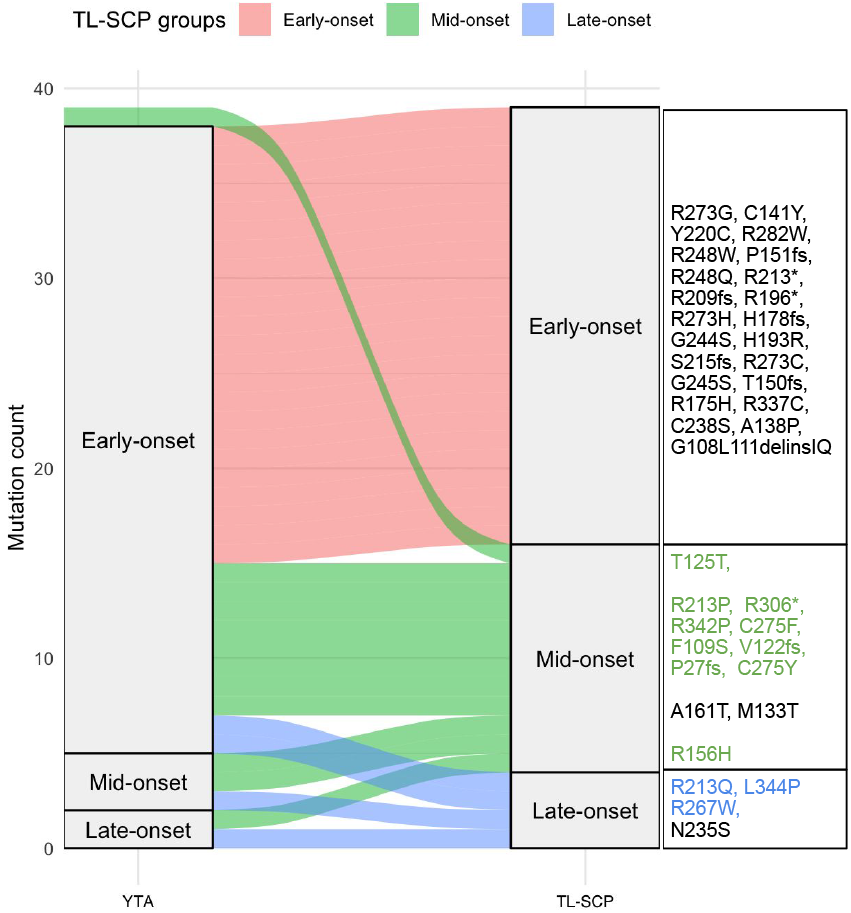
Alluvial plot of *TP53* mutation annotations (YTA vs. TL-SCP). The right-hand text boxes list individual mutations in each TL-SCP group. Mutations highlighted in color indicate those with discrepant annotations between YTA and TL-SCP.

**Figure 6.**
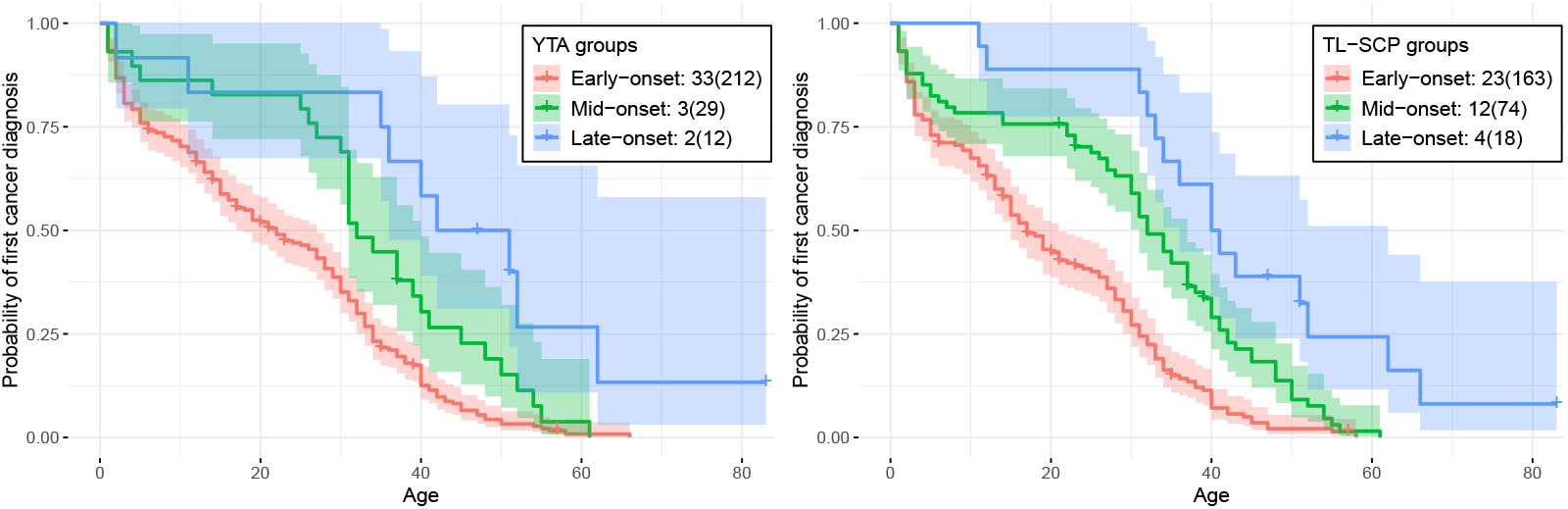
Kaplan–Meier survival curves (in age at first cancer diagnosis with 90% confidence intervals) of LFS patients stratified by the three mutation groups identified by YTA (left panel) and TL-SCP (right panel). Numbers in the legend indicate the number of mutations, with the corresponding number of patients in parentheses.

## 5 Discussion

Motivated by the pressing need for *TP53* mutation annotation in cancer biology, we introduced Survival-based Clustering of Predictors (SCP), a general framework for grouping individual features, such as single-nucleotide variants, by searching for homogeneity in their coefficients in Cox regression. SCP builds on a fusion-penalized Cox model and employs a distance-to-set penalty to facilitate parameter tuning and enhance clustering performance. To address common data limitations in survival analysis, we further developed TL-SCP, a transfer learning-adapted extension that leverages coefficient ranking information from a source cohort. This novel ranking-based transfer assumption is less restrictive than the magnitude-based assumptions in conventional transfer learning, thereby making TL-SCP more broadly applicable in practice. Comprehensive simulation studies demonstrated the superior performance of TL-SCP, as well as its ability to accommodate cohort heterogeneity and ranking inconsistency. When applied to a population database of individuals with germline *TP53* mutations, TL-SCP achieved more biologically and clinically meaningful *TP53* mutations clusters, compared to the state-of-art in vitro experimental functional assay results. All functions and related analyses are publicly available at https://github.com/Xiaoqian-Liu/TL-SCP.

We foresee many potential directions for future investigation. First, while TL-SCP and SCP currently use the classic Cox model, alternative survival models could enable more advanced analyses and potentially improve clustering performance. Second, the ranking-transfer strategy may be extended to other statistical learning contexts to enhance target-task performance. In addition, more principled approaches for integrating ranking information from the source could be developed to replace the current ad-hoc weighted rank averaging. Third, there are many opportunities to apply TL-SCP/SCP to biomedical research questions. While this study focused on a pan-cancer analysis with age at first cancer diagnosis as the outcome, the method or its variants could be adapted for cancer type–specific *TP53* mutation annotation, where data limitations are more pronounced. In such contexts, integrating transfer learning with domain knowledge holds strong potential to address these challenges. We can also apply TL-SCP/SCP to group effects of somatic mutations in *TP53* or other frequently mutated cancer genes such as *BRCA1/2*. With the huge amount of genomic data already available for cancer patients, TL-SCP and SCP hold strong promise for identifying clustered genetic features that may explain time-to-event outcomes such as progression-free or overall survival, creating new opportunities in data mining to expedite discoveries for improved prevention and treatment strategies in cancer and other diseases.

## Supplementary Materials

Supplementary Materials to

